# Complete mitochondrial genomes of ancyromonads provide clues for the gene content and genome structures of ancestral mitochondria

**DOI:** 10.1101/2025.03.26.645259

**Authors:** Ryo Harada, Takashi Shiratori, Akinori Yabuki, Yuji Inagaki, Andrew J. Roger, Ryoma Kamikawa

## Abstract

Mitochondria of eukaryotic cells are direct descendants of an endosymbiotic bacterium related to Alphaproteobacteria. These organelles retain their own genomes, which are highly reduced and divergent when compared to those of their bacterial relatives. To better understand the trajectory of mitochondrial genome evolution from the last eukaryotic common ancestor (LECA) to extant species, mitochondrial genome sequences from phylogenetically diverse lineages of eukaryotes – particularly protists – are essential. For this reason, we focused on the mitochondrial genomes of Ancyromonadida, an independent and understudied protist lineage in the eukaryote tree of life. Here we report the mitochondrial genomes from three Ancyromonadida: *Ancyromonas sigmoides*, *Nutomonas longa*, and *Fabomonas tropica*. Our analyses reveal that these mitochondrial genomes are circularly mapping molecules that feature protein-coding inverted repeats. This inverted repeat structure has been observed in other mitochondrial genomes but is patchily distributed over the tree of eukaryotes. Ancyromonad mitochondrial genomes possess several protein-coding genes which have not been detected from any other mitochondrial genomes of eukaryotes sequenced to date, thereby extending the known mitochondrial gene repertoire of ancestral eukaryotes, including LECA. These findings significantly expand our understanding of mitochondrial genome diversity across eukaryotes, shedding light on the early phases of mitochondrial genome evolution in eukaryotes.

## TEXT

Mitochondria are double-membrane bounded organelles that are best known as the ATP-producing “powerhouses” of extant eukaryotic cells. They are the direct descendants of endosymbiotic bacteria related to Alphaproteobacteria that were integrated into host cells of archaeal origin during eukaryogenesis (Richards et al., 2024; Roger et al., 2017; Vosseberg et al., 2024). Except for several anaerobic groups (Abrahamsen et al., 2004; Leger et al., 2017; Williams et al., 2002), the mitochondria of most eukaryotic species retain their own genomes derived from their endosymbiotic bacterial ancestor. However, the vast majority of the genes of the original endosymbiont were either lost if they were no longer necessary for the host– endosymbiont partnership or transferred to the nuclear genome during genetic and functional integration between the endosymbiont and the host. The end result are mitochondrial genomes of extant species that are streamlined in size and gene content compared to their bacterial relatives (Ettema and Andersson, 2009). Importantly, even though some of this genomic streamlining occurred during eukaryogenesis, it has continued to occur after the diversification of the major extant eukaryotic lineages from the last eukaryotic common ancestor (LECA), yielding a considerable variation in gene content even among mitochondrial genomes of relatively closely related protist species (Eglit et al., 2024; Kamikawa et al., 2016; Moreira et al., 2024; Yazaki et al., 2022).

The most gene-rich mitochondrial genomes currently known are those of the discobid group Jakobida with 65–100 kb in length, which carry 61–66 protein genes and 30–34 RNA genes (Burger et al., 2013). The most gene-rich mitochondrial genome outside Jakobida so far was found in the CRuMs, representatives of which have 53–63 kb-long mitochondrial genomes carrying 50–62 protein genes and ∼30 RNA-encoding genes (Kamikawa et al., 2016; Moreira et al., 2024). At the other end of the spectrum, the smallest mitochondrial genomes are found in dinoflagellates, apicomplexan parasites, and their photosynthetic close relatives, that retain only two or three protein genes and fragmented rRNA genes (Oborník and Lukeš, 2015). The mitochondrial genomes of apicomplexan parasites are also the smallest in size (6–7 kb; Feagin, 2000). In addition to their coding capacity, mitochondrial genomes show substantial diversity in structure among species (Burger et al., 2003; Gray et al., 1998). The vast majority of aerobic eukaryotes possess circularly- or linearly-mapping mitochondrial genomes. However, mitochondrial genomes of dinoflagellates are composed of a variety of linear DNA fragments generated by recombination (Kamikawa et al., 2007; Norman and Gray, 2001; Slamovits et al., 2007). The mitochondria of some Euglenozoa also have unusual features, including gene fragmentation where each gene fragment is located within small circular DNAs (Dobáková et al., 2015; Kaur et al., 2020; Spencer and Gray, 2011; Verner et al., 2015). Although somewhat unusual, inverted repeats including protein-coding genes (protein-coding inverted repeats; PCIR) are present in the mitochondrial genomes of a variety of distantly-related protists including *Malawimonas jakobiformis* (NC_002553), *Proteromonas lacertae* (Pérez-Brocal et al., 2010), *Palpitomonas bilix* (Nishimura et al., 2016), and *Acavomonas peruviana* (Janouškovec et al., 2013; Tikhonenkov et al., 2014). To accurately infer the gene content and structure of the mitochondrial genome of LECA and the subsequent evolutionary ‘paths’ from LECA to various extant species, we need broad sampling of mitochondrial genomes of phylogenetically diverse eukaryotes, particularly those from deep-branching lineages.

One important understudied ‘deep branching’ protist lineage is the Ancyromonadida, a group of flagellates that are found in diverse aquatic and soil environments (Heiss et al., 2010; Yubuki et al., 2023). Members of Ancyromonadida are heterotrophic protists, characterized by two flagella, round-to-bean-shaped cells and a very small cell size of approximately 4 ± 0.5 µm in length. Electron microscopic analyses showed that these cells have a pellicle under the cell membrane and extrusomes (Heiss et al., 2010; Yubuki et al., 2023). Recent phylogenomic analyses have not settled on a robust placement of the Ancyromonadida in the eukaryote tree. Recent nuclear gene based analyses position Ancyromonadida as a lineage closely related to Malawimonadida and CRuMs (Eglit et al., 2024; Harada et al., 2024; Torruella et al., 2025; Yazaki et al., 2025), which, together with Amorphea comprised of Opisthokonta, Apusomonadida, Breviatea, and Amoebozoa, are proposed to form the huge assemblage called Opimoda (Derelle et al., 2015). The phylogenetic group including Ancyromonadida in Opimoda is referred to as “Opimoda+” (Williamson et al., 2025). The latest mitochondrial protein-based phylogenomic analyses placed the eukaryotic root position between Opimoda+ and Diphoda+, the latter of which comprises Diaphoretickes and Discoba (Williamson et al., 2025).

Recently, Gastineau et al. (2023) reported two incomplete linear mitochondrial genome fragments of Ancyromonadida: one from *Ancyromonas sigmoides* strain 1C2 and another from metagenomic samples. Given the incomplete mitochondrial genome fragments of *A. sigmoides* and lack of any mitochondrial genome sequence in other ancyromonad lineages, diversity and evolution of the mitochondrial genome structures and gene contents in Ancyromonadida remain poorly understood. To expand our understanding of mitochondrial genome diversity in eukaryotes, we have characterized the complete mitochondrial genome sequences of representatives of three ancyromonad species: *A. sigmoides*, *Nutomonas longa*, and two strains of *Fabomonas tropica*. Our analyses reveal that these Ancyromonadida mitochondrial genomes possess protein-coding inverted repeats (PCIRs) and some ribosomal protein genes that have not previously been found in mitochondrial genomes. These findings update our understanding of diversity in mitochondrial genome structure and gene repertoire across the eukaryote tree, affording deeper insights into the evolutionary history of these organelles.

## MATERIALS AND METHODS

### Culturing, sequencing, and assembling

Cells of *A. sigmoides* B-70, *N. longa* NCFW, and *F. tropica* NYK3C were provided by Dr. Aaron A. Heiss (Glücksman et al., 2013; Heiss et al., 2010). The protist DNA extracted with Plant DNA Extraction Kit (Jena Biosciences) were subjected to the library preparation with the Illumina TruSeq Nano DNA Library Prep Kit for 350 bp inserts according to the manufacturer’s instructions and sequenced with the HiSeq 2500 (Illumina), resulting in 46.0 million, 44.1 million, and 44.5 million paired-end reads for *A. sigmoides* B-70, *N. longa* NCFW, and *F. tropica* NYK3C, respectively. Adapter trimming was conducted using Cutadapt version 1.1 (Martin, 2011) as the default setting. Quality filtering was conducted with Trimmomatic v0.32 (Bolger et al., 2014), setting the SLIDINGWINDOW option to 20:20 and keeping the minimum length of 50 nt. The resulting reads were assembled using Velvet version 1.2.08. (Zerbino and Birney, 2008), setting input reads, expected coverage, and hash length as the short paired-end reads, “auto,” and 35-55, respectively. By BLAST-based homology searches using *Reclinomonas americana* mitochondrial proteins (NC_001823.1), mitochondrial genome-derived contigs were detected as fragments of mitochondrial DNAs. Taking overlapping sequences at the 5ʹ and 3ʹ ends and read coverage into consideration, the contigs were manually assembled into single circularly mapping DNAs. Quality-filtered reads were then mapped onto the DNAs by HISAT2 (Kim et al., 2019) using default settings to confirm that the manual assemblies were correct (Fig. S1).

Cells of *F. tropica* SRT902 were cultivated as previously described (Harada et al., 2024). The whole DNA extracted by the phenol-chloroform method was subjected to the library preparation with the VAHTS Universal DNA Library Prep Kit for Illumina V3 ND607 and sequenced with the HiSeq X (Illumina), resulting in 112 million paired-end reads. Library construction and sequencing were conducted at a biotech company (AZENTA Japan Corp., Tokyo, Japan). The raw reads of *F. tropica* SRT902 were trimmed with fastp v0.19.7 (Chen et al., 2018) with the ‘-q 20 -u 80’ options and then assembled with SPAdes v3.15.3 (Nurk et al., 2017) using the ‘-meta -k 13,21,29,37,45,53,61,69,77,85,93,101,109,117,125’ options. Mitochondrial contigs were detected by BLAST-based homology search using *Jakoba libera* mitochondrial proteins (NC_021127.1). The contigs were manually assembled into single mapping DNA. Quality-filtered reads were then mapped onto the assembly by HISAT2 (Kim et al., 2019) as the default settings to confirm that the manual assembly was correct.

We also sequenced the mitochondrial genome of the CRuM species *Mantamonas sphyraenae* SRT306. However, as we were completing our analyses, the same organisms’ mtDNA was published by Moreira et al. (2024). Nevertheless, we briefly describe our methods here. The *M. sphyraenae* cells were provided by Dr. Aaron A. Heiss. The *M. sphyraenae* DNA extracted with Plant DNA Extraction Kit (Jena BioSciences) was subjected to the library preparation with the Illumina TruSeq Nano DNA Library Prep Kit for 350 bp insert, according to the manufacturer’s instruction and sequenced with the Illumina HiSeq 2500, resulting in 47.6 million paired-end reads. Adapter trimming was conducted using Cutadapt version 1.1 (Martin, 2011) as the default setting. Quality filtering was conducted with Trimmomatic v0.32 (Bolger et al., 2014), setting the SLIDINGWINDOW option as 20:20 and keeping the minimum length of 50 nt. The resulting reads were assembled using Velvet version 1.2.08. (Zerbino and Birney, 2008), setting input reads, expected coverage, and hash length as the short paired-end reads, “auto,” and 65, respectively. A mitochondrial DNA survey was conducted with BLAST-based homology search (Camacho et al., 2009) using *R. americana* mitochondrial proteins as described above. Out of the resultant contigs, we found one contig with 53,109 nt and 26.52 GC% that formed a circularly mapping molecule of 53,051 bp with 58 nt-overlapping sequences at the 5ʹ and 3ʹ ends. Since the mitochondrial genome sequence of *M. sphyraenae* has already been reported by Moreira et al. (2024), we only briefly mention this genome in the following sections and do not discuss it in detail.

### Gene annotation

The mitochondrial genome sequences of four Ancyromonadida strains and *M. sphyraenae* were annotated using the MFannot (Lang et al., 2023) web server with the standard mitochondrial genetic code (NCBI’s translation table 1). Infernal v1.1.5 (Nawrocki and Eddy, 2013) and covariance model of mt-tmRNA (RF02544) were used to search for transfer-messenger RNA (tmRNA). Functional annotation of each open reading frame (ORF) assigned by MFannot was confirmed by PSI-BLAST against clustered_nr as of September 2024 (Altschul et al., 1997). If annotations of genes by MFannot and PSI-BLAST were inconsistent with each other, they were further investigated and their annotation was treated as provisional. ORFs not confidently annotated in MFannot and PSI-BLAST with E-values below 1e-4 were subjected to the following, more detailed annotation. For functionally unknown ORFs, we performed conserved domain searches using InterProScan v5.69 (Jones et al., 2014). The 3D structure of each ORF was predicted by the AlphaFold3 web server (https://alphafoldserver.com) v2024.08.19 (Abramson et al., 2024) and the predicted tertiary structures were used as queries in a similarity search using the Foldseek web server (https://search.foldseek.com) as of September 2024 (van Kempen et al., 2024) with an E-value cutoff of 1e-4.

Furthermore, an all-against-all BLAST analysis was performed among the four Ancyromonadida mitochondrial genomes. We focused on *atp4*, *sdh3*, and *tatC* in *A. sigmoides*, which were absent in the other three Ancyromonadida mitochondrial genomes. The tertiary structures of these three proteins were predicted using AlphaFold3 and compared with the structures of functionally unknown ORFs in the other three mitochondrial genomes using TM-align (Zhang and Skolnick, 2005) with a TM-score cutoff of 0.5 to search for potential homologues. Additionally, we assessed transmembrane (TM) domains in functionally unknown ORFs using TMHMM2.0 (Krogh et al., 2001), DeepTMHMM1.0 (Hallgren et al., 2022), and Phobius1.01 (Käll et al., 2004). Genome maps were drawn by OGDRAW (Greiner et al., 2019) and edited manually.

With the above-mentioned tools, we reinvestigated the gene contents of two previously reported Ancyromonadida mitochondrial genomes, Ancyromonadida sp. SAT37 (PALT01000012.1) and *A. sigmoides* 1C2 annotated in Gastineau et al. (2023). We found one gene of Ancyromonadida sp. SAT37 was annotated as *rps10* in Gastineau et al. (2023), which was not detected in the mitochondrial genomes of either *A. sigmoides* 1C2 (Gastineau et al., 2023) or B-70 (this study). No tools we employed identified it as *rps10*; therefore, we treat Ancyromonadida sp. SAT37 as a mitochondrial genome lacking *rps10*.

## RESULTS

### Mitochondrial genome structure

The complete mitochondrial genome of *A. sigmoides* B-70 was determined to be a circularly-mapping molecule, with a genome size of 48,960 bp (Fig. 1; Table 1). This genome exhibited a G+C content of 34% and coding regions accounted for 90% of the genome sequence. Mitochondrial small subunit rRNA (SSU rRNA) gene in *A. sigmoides* B-70 showed 99.8% identity to the previously sequenced strain 1C2. Despite this high degree of similarity, the mitochondrial genome of B-70 was found to be 7,071 bp larger than the published mitochondrial sequence from strain 1C2, likely due to the incompleteness of the latter genome (Gastineau et al., 2023). Notably, the B-70 mitochondrial genome contains PCIRs spanning 7,108 bp, which nearly matches the size discrepancy observed in strain 1C2, supporting the notion that the latter may be missing this sequence. In fact, the gene content and synteny of the mitochondrial genomes of B-70 and 1C2 were identical except for presence/absence of the PCIR regions. In the raw read mapping analysis, if the PCIRs are indeed repeat regions, the read coverage on the PCIRs would be approximately two times higher than that of single-copy regions. Consistent with this prediction, the coverage of the PCIR region and the single-copy regions were 149.77× and 75.21×, respectively in B-70 (Table 1).

**FIGURE 1.**
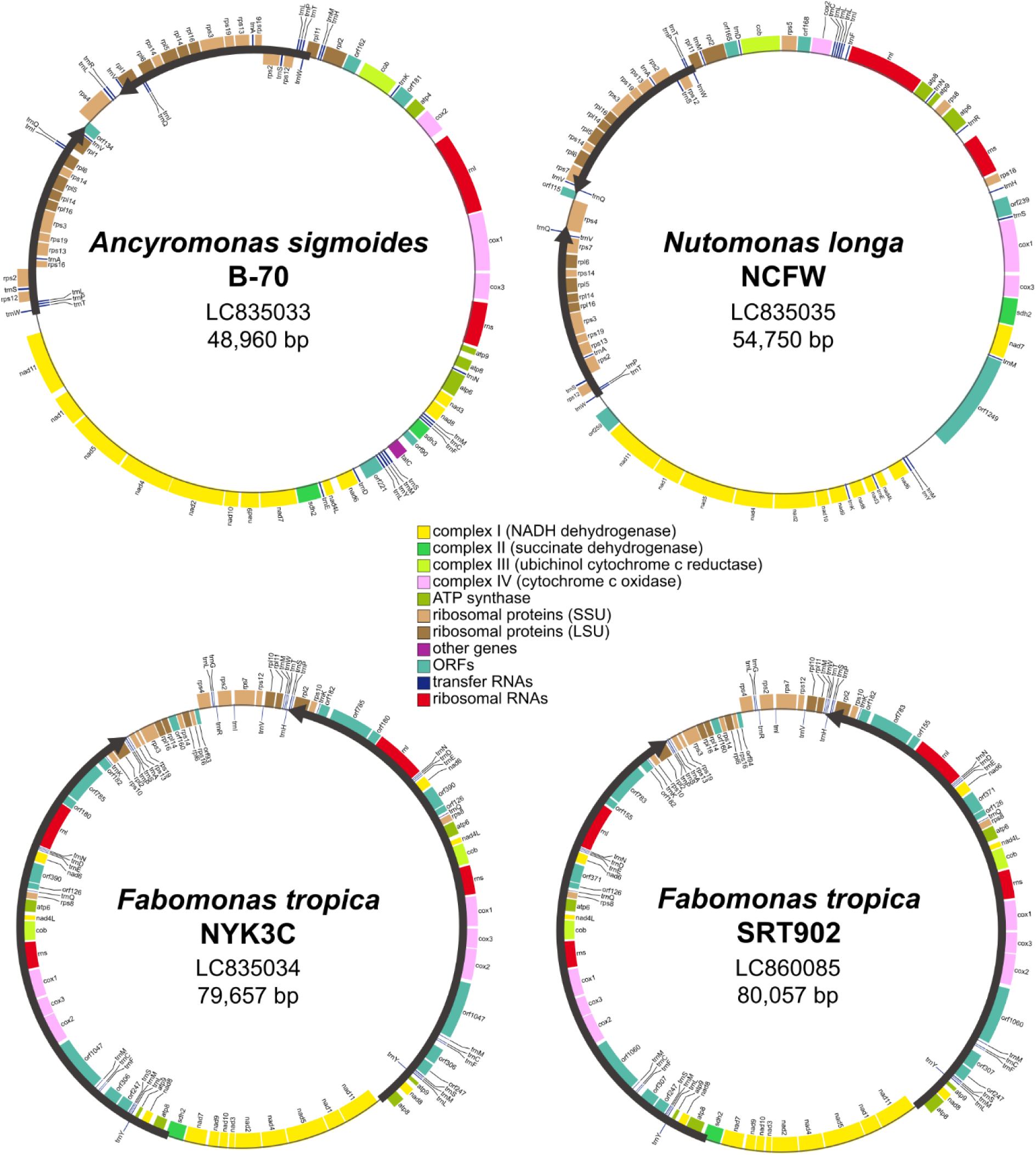
The complete genome maps of four Ancyromonadida species mitochondrial genomes. Coding regions are colored based on their function (see colour key in figure). Black bold arrows indicate inverted repeats. Each map was drawn by OGDRAW and edited manually.

**TABLE. 1.** Characteristics of mitochondrial genomes from Ancyromonadida and phylogenetically related candidates.

|  | accession | genome size (bp) | GC (%) | ORFs | coding ratio (%) | protein-coding inverted repeats (bp) | coverage of inverted repeats/single-copy region |
| --- | --- | --- | --- | --- | --- | --- | --- |
| *Ancyromona sigmoides* B-70 | LC835033 | 48,960 | 34.18 | 55 | 90.31 | 7,108 | 149.77×/75.21× |
| *Nutomonas longa* NCFW | LC835035 | 54,750 | 34.45 | 54 | 88.96 | 7,052 | 20.46×/10.06× |
| *Fabomonas tropica* NYK3C | LC835034 | 79,657 | 25.28 | 68 | 91.22 | 28,695 | 449.11×/259.30× |
| *Fabomonas tropica* SRT902 | LC860085 | 80,057 | 25.39 | 68 | 91.46 | 28,898 | 681.40×/381.73× |
| *Malawimonas jakobiformis* | AF295546 | 47,328 | 26.11 | 49 | 91.13 | 9,100 | - |
| *Malawimonas californiana* | KP165387 | 36,792 | 26.79 | 39 | 89.72 | 608 | - |
| *Diphylleia rotans* | NC_029886 | 62,563 | 34.40 | 58 | 68.80 | 0 | - |
| *Mantamonas sphyraenae* | LC842150 | 53,047 | 26.54 | 63 | 90.98 | 0 | - |

The *A. sigmoides* B-70 mitochondrial genome carries two rRNA genes and 24 tRNA genes: the annotation program MFannot did not detect a conserved 5S rRNA gene. The mitochondrial genome encodes 38 proteins with known functions when counting duplicated genes as one (Fig. 1), and no introns were detected in any of the detected ORFs. The mitochondrial gene content of *A. sigmoides* B-70 is identical to that of strain 1C2. Besides possessing genes encoding the eight small subunit ribosomal proteins (rps) and seven large subunit ribosomal proteins (rpl), it retains genes encoding subunits of complex I (*nad1-4, 4L, and 5-11*), complex II (*sdh2* and *sdh3*), complex III (*cob*), complex IV (*cox1-3*), ATP synthase complex (*atp4, 6, 8, and 9*), and the twin arginine translocator (*tatC*). The PCIRs encode 9 tRNA (*trnA, I, L, P, Q, S, T, V, and W*), 7 rps (*rps2, 3, 12, 13, 14, 16, and 19*), and 5 rpl (*rpl1, 5, 6, 14, and 16*) genes.

The mitochondrial genome of *N. longa* NCFW was also a circularly-mapping molecule, with a genome size of 54,750 bp (Fig. 1; Table 1). The genome retains two rRNA genes and 24 tRNA genes. The *N. longa* mitochondrial genome encodes 37 proteins with known functions. Introns were not detected in any ORFs. Besides the 17 ribosomal proteins comprised of 11 rps and 6 rpl, the *N. longa* mitochondrial genomes has genes encoding subunits of complex I (*nad1-4, 4L, and 5-11*), complex II (*sdh2*), complex III (*cob*), complex IV (*cox1-3*), and ATP synthase (*atp6, 8, and 9*), but lacks *sdh3*, *atp4* and *tatC* that are present in the mitochondrial genome of *A. sigmoides*. Seven tRNA (*trnA, P, Q, S, T, V, and W*), 7 rps (*rps2, 3, 7, 12, 13, 14, and 19*), and 4 rpl (*rpl5, 6, 14, and 16*) genes were found to be present within the PCIRs of *N. longa*.

The sequenced mitochondrial genomes of two strains of *F. tropica*, NYK3C and SRT902, were determined as circularly-mapping molecules with genome sizes of 79,657 bp and 80,057 bp, respectively (Fig. 1; Table 1). Mitochondrial SSU rRNA genes of the two *F. tropica* strains shared 98% identity with each other. Both genomes exhibited 25% G+C content and 91% coding capacity. The two *F. tropica* strains exhibit larger PCIRs, measuring 28,695 bp and 28,898 bp, than those in the other Ancyromonadida species sequenced in this study (Table 1). The gene content and synteny of the two *F. tropica* strains are identical to each other. They encode two rRNA genes and 24 tRNA genes. The *F. tropica* mitochondrial genomes encode 37 proteins. Introns were not detected in any ORFs. Besides the ribosomal proteins, the protein gene contents are identical among strains of *N. longa* and *F. tropica*. Their gene contents differed only in presence or absence of *rps5*, *rps10*, *rpl5*, and *rpl10*. In the PCIRs, 14 tRNA (*trnC, D, E, F, K, L, M, N, P, Q, S, and Y*) including two copies each of *trnM* and *trnS*, 2 rps (*rps8 and 10*), 1 rpl (*rpl2*), 2 rRNA (*rnl and rns*), 3 nad (*nad4L, 6, and 8*), *cob*, 3 cox (*cox1-3*), and 3 atp (*atp6, 8, and 9*) genes as well as 8 unidentified ORFs (orf126, 180, 182, 247, 306, 390, 785, and 1047) are present.

The mitochondrial gene synteny in Ancyromonadida is conserved. Specifically, there are conserved operon-like gene clusters with adjacent genes encoding functionally related subunits (Fig. 1, 2A, 2B). For example, we found that *nad11-nad1-nad5-nad4-nad2*, *nad10-nad9*, *rps13-rps19-rps3-rpl16-rpl14*, *cox1-cox3*, and *nad7-sdh2* are conserved among all the genomes analyzed in this study. Collectively, the four genomes sequenced here represent the first complete mitochondrial genomes from Ancyromonadida.

**FIGURE 2.**
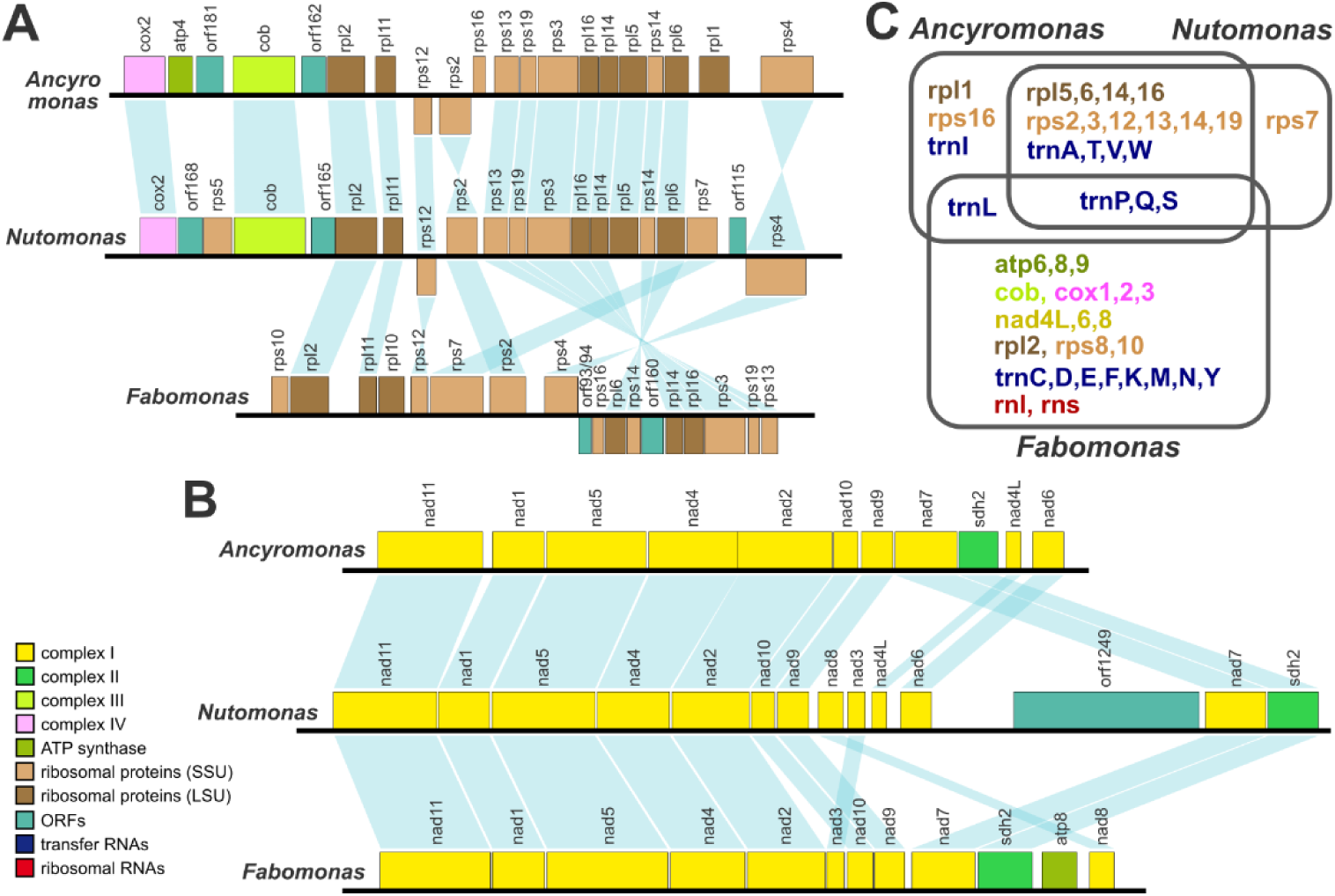
Conservation of synteny and gene repertoire of inverted repeats. Conservation of synteny around ribosomal genes (**A**) and complex I genes (**B**) is highlighted in blue band. tRNA genes are omitted in synteny analysis. (**C**) Gene repertoire of inverted repeats.

### Functionally uncharacterized ORFs

Out of all the functionally annotated genes with MFannot and/or PSI-BLAST with significant statistical reliability (E-value < 1e-4), only *rps5* of *N. longa* and *rpl10* of *F. tropica* strains were annotated by more detailed analyses using InterProScan. In addition to this prediction, the amino acid sequences of the ORFs that were not confidently annotated in MFannot and PSI-BLAST were subjected to an AlphaFold/FoldSeek analysis, which confirmed that the tertiary structures predicted by AlphaFold3 for the proteins encoded by *N. longa* orf198 (Nuto01.10) and *F. tropica* NYK3C/SRT902 orf188/orf189 (Fabo01.15/SRT902.15) hit the tertiary structures of Rps5 and Rpl10 with E-values of 3.8e-5 and 6.1e-6/6.2e-5, respectively (Table S1). We also compared the tertiary structures of *N. longa* Rps5 and *F. tropica* Rpl10 with those determined by cryo-electron microscopy of Human mitochondrial ribosomes (Fig. 3). *N. longa* Rps5 was aligned with the root mean squared deviations (RMSDs) of 3.86 and 1.94 in the N- and C-terminal regions, respectively, and *F. tropica* Rpl10 was aligned with RMSD of 2.58. To date, this is the only known case of Rps5 being encoded on a mitochondrial genome. Based on Foldseek results (Table S1), *F. tropica* orf182 (Fabo01.11) seems likely that it might encode Rps5, although no significant sequence similarity between translated orf182 and Rps5 was detected by using all the above-mentioned tools. Using amino acid sequences of *N. longa* Rps5 and *F. tropica* Rpl10 as queries, homology-based surveys also did not detect any homologs of these proteins in the mitochondrial genomes of three Ancyromonadida species.

**FIGURE 3.**
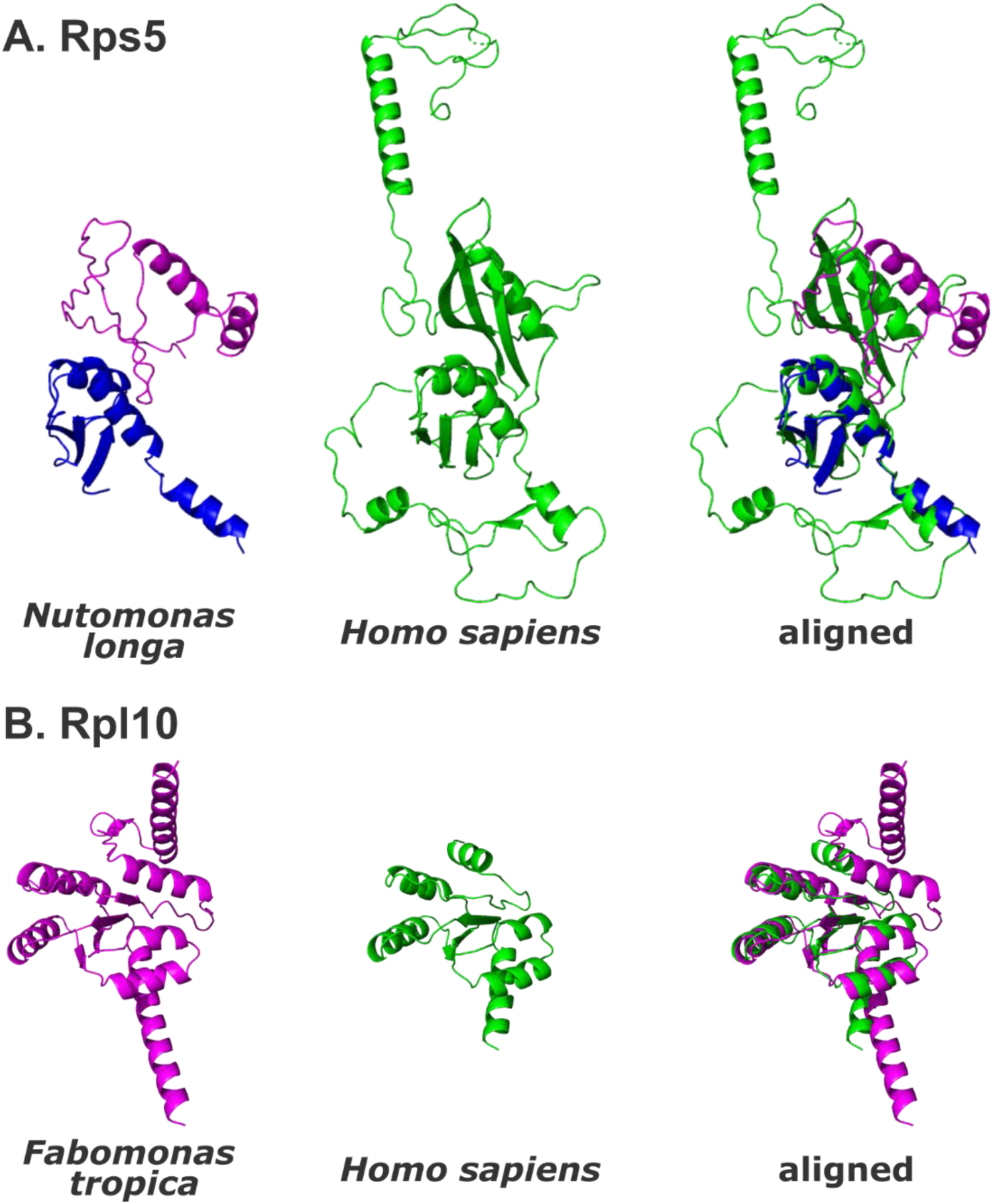
Structural comparison of Rps5 and Rpl10. (**A**) The N-terminal (1–77 amino acid residues) and C-terminal (119–198 amino acid residues) regions of *Nutomonas longa* Rps5 are shown in magenta and blue, respectively. The green color corresponds to the experimentally determined structure of *Homo sapiens* Rps5 (126–430 amino acid residues of chain AD in pdb_00003j9m). (**B**) *Fabomonas tropica* NYK3C Rpl10 and *H. sapiens* Rpl10 (77–197 amino acid residues of chain I in pdb_00003j9m) are shown in magenta and green, respectively. Structures were aligned by TM-align.

MFannot annotated the C-terminal region of *N. longa* Orf1249 with 900 amino acid residues as Sdh3, a subunit of mitochondrial complex II with a typical length of 100-200 amino acids. However, our subsequent analyses with PSI-BLAST, InterProScan, and AlphaFold/Foldseek did not confirm the similarity to Sdh3, leaving Orf1249 functionally unidentified. The predicted 3D structure of Orf1249 showed alpha helices aligned along the same axial direction (Fig. 4). Three *in silico* tools, DeepTMHMM, TMHMM2 and Phobius, also predicted some of the helices as TM domains. Since Sdh3 is known to be a membrane protein containing a TM domain as part of Complex II, it is possible, but not likely, that the initial MFannot assignment of Orf1249 as Sdh3 could be correct. The tertiary structures of *A. sigmoides* Sdh3 and *N. longa* Orf1249 were aligned using TM-align, but the RMSD was 5.22, indicating no significant structural similarity. Another possibility is that because Orf1249 is much longer than typical Sdh3, it could correspond to Sdh3 fused with another protein or other proteins. Gene fusions are observed in the mitochondrial genomes of Opimoda+, e.g. *cox1/cox3* in *Gefionella okellyi* (a member of Malawimonadida) and *cox1/cox2* in *Acanthamoeba castellanii* (a member of Amoebozoa) (Lonergan and Gray, 1996; Valach et al., 2014). However, due to lack of definitive sequence and structural similarity evidence, precise annotation of orf1249 remains difficult.

**FIGURE 4.**
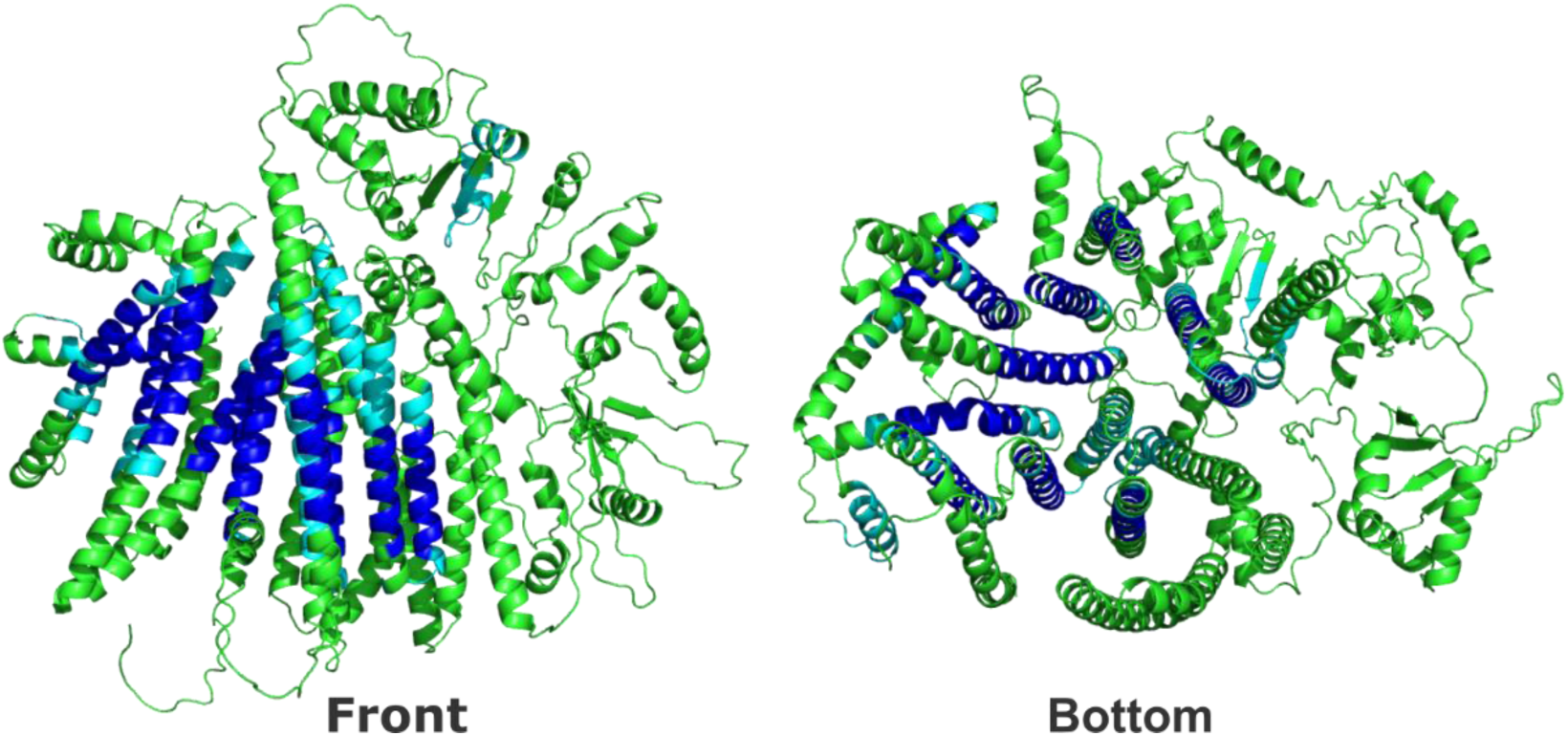
Predicted 3D structure and TM helices of *Nutomonas longa* Orf1249. The structure of *N. longa* Orf1249 predicted by AlphaFold3 is shown in the front view and bottom view on the left and right, respectively. Transmembrane (TM) helices predicted by three *in silico* tools are shown in blue. TM helices predicted by at least one of three tools are shown in cyan.

In any case, genes encoding functionally annotated proteins larger than 1,000 amino acids – e.g., *rpoB* and *rpoC*, which are conserved in Jakobida species – are rare in mitochondrial genomes. It is interesting that the mitochondrial genomes of Ancyromonadida, which have relatively high coding density, retain large ORFs of unknown function. At present, we have no clues to predict functions encoded in the other uncharacterized ORFs of the ancyromonad mitochondrial genomes, including the large ORFs in *F. tropica*.

## DISCUSSION

We determined the complete genome sequences of mitochondria in four ancyromonad strains from three species, *A. sigmoides*, *N. longa*, and *F. tropica*. As in some other mitochondrial genomes found in diverse eukaryotes, the mitochondrial genomes of ancyromonads are A+T rich, circularly-mapping molecules that are densely packed by coding regions. Almost all the genes found in the mitochondrial genomes of Ancyromonadida are those that have previously been reported. However, we report a *rps5* gene that has never been found in any previously published mitochondrial genomes, as well as a *rpl1* gene that has never been found in the previously published Opimoda+ mitochondrial genomes. In addition to the functionally annotated genes, the Ancyromonadida mitochondrial genomes carry many ORFs that do not appear to have significant similarity to functionally annotated proteins in any databases. Unfortunately, even predicted tertiary structure-based analyses for all the ORF provides no clue to estimate functions of their encoded proteins. Nevertheless, it is intriguing that 15 out of the 34 ORFs were predicted to have one or more TM domains (Table S1), suggesting that they might encode membrane-localized proteins such as transporters of metabolites. There are three possibilities for each of the functionally unannotated ORFs: i) they might be non-coding sequences, ii) they might be newly produced genes that possess no homologues in other organisms, or iii) they might be divergent homologues of functionally known proteins. The high coding density of these genomes suggests that i) is unlikely. In any case, with the expansion of available mitochondrial genome sequence data and the improvement of homology detection methods, annotation of these ORFs might be possible in future. Alternatively, if these are truly “new genes,” biochemical experiments will be needed to determine their functions.

### Ancestral genome structures and the evolution of mitochondria

All of the four complete mitochondrial genomes of Ancyromonadida have PCIRs. In *A. sigmoides* and *N. longa*, these PCIRs are 7,108 bp and 7,052 bp, respectively, sharing similar gene repertoires and orders (Fig. 1 arrows, 2C). On the other hand, the two *F. tropica* strains, NYK3C and SRT902, have much larger PCIRs that are 28,695 bp and 28,898 bp long, respectively, with gene contents and orders identical to each other. However, almost all the genes on the PCIRs are not shared between *Ancyromonas*/*Nutomonas* and *Fabomonas*, and only three tRNA genes are shared among them (Fig. 2C). A recent phylogenomic analysis suggested that *A. sigmoides* and *N. longa* are more closely related to each other than to *F. tropica* (Brown et al., 2018; Gastineau et al., 2023; Harada et al., 2024; Torruella et al., 2025; Williamson et al., 2025; Yazaki et al., 2025), which is consistent with the patterns of similarities/differences of the PCIRs of these taxa. The close relationship and the similarity of PCIRs between *A. sigmoides* and *N. longa* suggest the last common ancestor of the two genera possessed PCIRs with a similar gene repertoire and gene order. Nevertheless, the stark differences in size and gene arrangement between PCIRs of *A. sigmoides*/*N. longa* and *F. tropica* strains make it difficult to infer ancestral features of PCIRs in acyromonads. As mentioned above, the PCIRs might have been present in the mitochondrial genome of the last common ancestor of Ancyromonadida followed by vertical inheritance to the extant species, and replacements of genes in the repeats. Alternatively, it is possible that PCIRs might have emerged independently after divergence in the *A. sigmoides*/*N. longa* lineage and in the *F. tropica* lineage. Currently it is unclear what mechanisms are underpinning the conversions between PCIRs-containing and PCIR-lacking mitochondrial genomes and how ‘easy’ these transitions are. It is known that plastid genomes in some photosynthetic algae have lost either copy of inverted repeats containing rRNA operons multiple times in evolution (Choi et al., 2019; Kamikawa et al., 2015; Kayama et al., 2020; Krämer et al., 2024; Matsuo et al., 2022; Palmer and Thompson, 1982), demonstrating that IRs can be lost.

More broadly, PCIRs have been reported in circularly-mapping mitochondrial genomes of phylogenetically diverse eukaryotes, including deep-branching lineages such as Malawimonadida, Nebulidia in Provora, *Meteora*, a heterolobosean strain BB2, and *Microheliella* (Eglit et al., 2024; Janouškovec et al., 2017; Tikhonenkov et al., 2022; Valach et al., 2014; Yang et al., 2017; Yazaki et al., 2022). It is also known that some linear mitochondrial genomes possess terminal PCIRs, such as *Palpitomonas*, *P. lacertae*, and *A. peruviana* (Janouškovec et al., 2013; Nishimura et al., 2016; Pérez-Brocal et al., 2010; Tikhonenkov et al., 2014). Analogous to the possibilities in Ancyromonadida discussed above, PCIRs might have emerged early in eukaryotic evolution followed by vertical inheritance, gene gains or losses within PCIRs, and multiple independent complete losses of PCIRs. Alternatively, PCIRs may have been acquired independently in multiple lineages. Since these are not mutually exclusive alternatives, a blend of both scenarios might have led to the punctate distribution of PCIRs observed across the eukaryotic tree. Since few species and lineages closely related to the above-mentioned eukaryotes that retain PCIRs have been sequenced, the extent to which PCIR-containing genome structures are conserved among closely related species remains uncertain, except for the Ancyromonadida. The detection of PCIRs in all the Ancyromonadida mitochondrial genomes completely sequenced in this study is intriguing in light of the not-yet fully resolved phylogenetic position of Ancyromonadida (Brown et al., 2018; Eglit et al., 2024; Harada et al., 2024; Torruella et al., 2025; Williamson et al., 2025; Yazaki et al., 2025); Representatives of Malawimonadida, one of the lineages potentially related to Ancyromonadida, also possess PCIRs in all their characterized mitochondrial genomes, although there is no PCIR gene shared by the two lineages. Nevertheless, if the sister group relationship between Ancyromonadida and Malawimonadida is true, the mitochondrial PCIRs found in Ancyromonadida and Malawimonadida might stem from vertical inheritance from their common ancestor followed by modification of their gene contents.

### Evolutionary history of mitochondrial gene repertoires

Our analyses of *A. sigmoides*, *N. longa*, and *F. tropica* have revealed variation in the mitochondrial gene repertoires of Ancyromonadida (Fig. 5). Whereas *rps5* is exclusively detected in the mitochondrial genome of *N. longa*, *rps10* and *rpl10* are exclusively detected in those of *F. tropica*. Similarly, *rpl1*, *atp4*, *sdh3*, and *tatC* are unique to *Ancyromonas* spp. In contrast, *rps7* and *rps8* are uniquely undetectable in the *Ancyromonas* mitochondrial genomes, whereas the lack of a detectable *rpl5* is unique to the *F. tropica* mitochondrial genomes. If we ignore the possibility of gene gain in mitochondrial genome evolution, then the last common ancestor of Ancyromonadida must have possessed all of the 43 protein-coding genes detected in the four mitochondrial genomes analyzed in this work. We believe it is reasonable to ignore gene gain by horizontal gene transfer in mitochondrial genomes, as only a few cases of such gain have been reported so far (He et al., 2016; Nishimura et al., 2020). During the diversification of the various lineages within Ancyromonadida, losses of genes or losses of detectable sequence homology must have happened differentially among species. Based on the closer relationship of *Ancyromonas* with *Nutomonas* than with *Fabomonas* (Brown et al., 2018; Gastineau et al., 2023; Harada et al., 2024; Torruella et al., 2025; Williamson et al., 2025; Yazaki et al., 2025), *rps5*, *rpl1*, *atp4*, *sdh3*, and *tatC* could have been lost (or diverged beyond recognition) multiple times independently in the evolution of Ancyromonadida.

**FIGURE 5.**
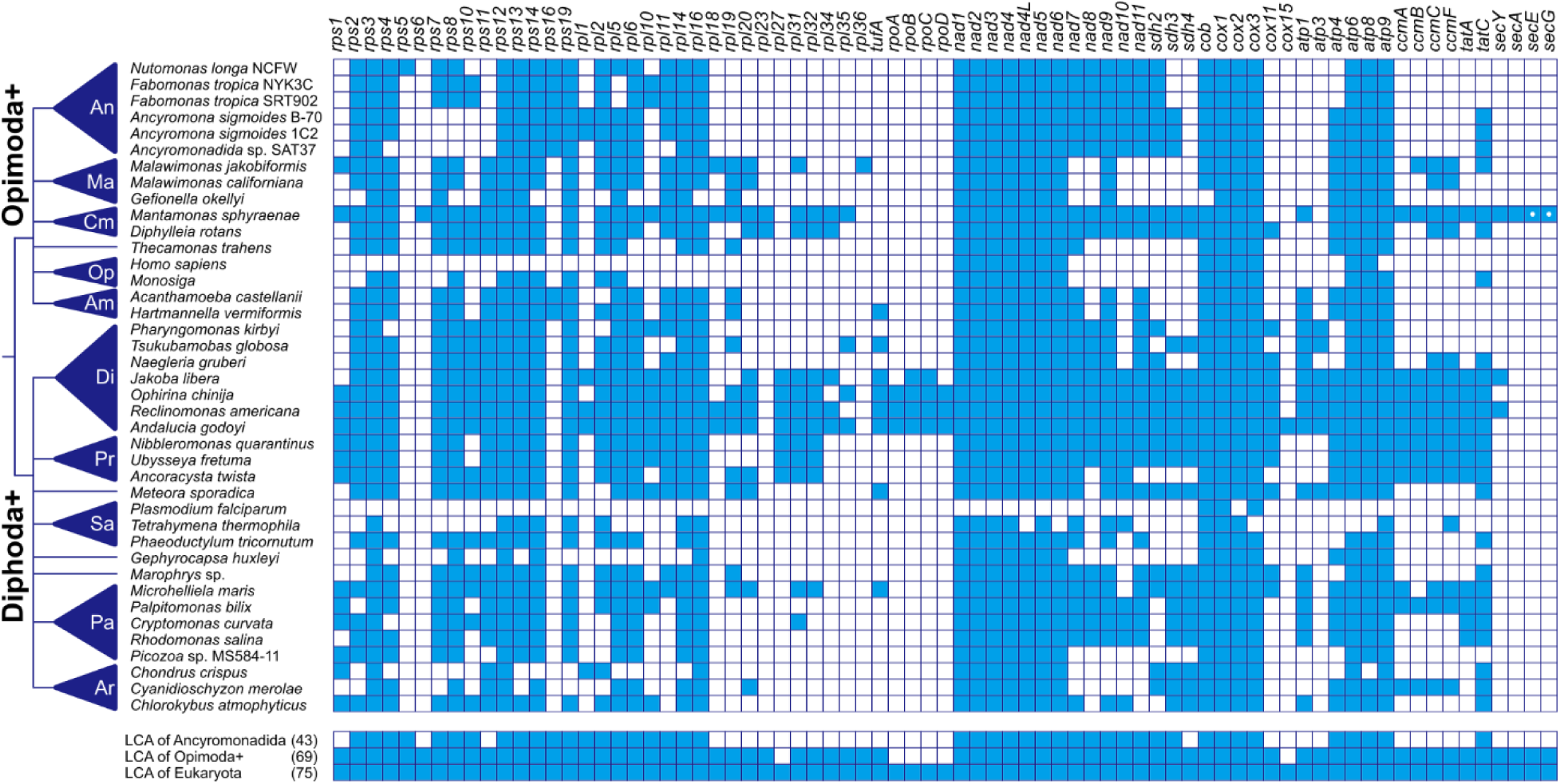
Repertoire of protein coding genes in mitochondrial genomes across the eukaryote tree of life. Presence and absence of corresponding genes of various eukaryotes is shown by blue and empty boxes, respectively. The figure is updated based on several previous studies (Eglit et al., 2024; Kamikawa et al., 2016; Moreira et al., 2024; Yazaki et al., 2022). Phylogenetic relationships of eukaryotes are based on Williamson et al., (2025). The predicted gene contents of the last common ancestor (LCA) of Opimoda+, LCA of Ancyromonadida, and LCA of Eukaryota are shown at bottom, and numbers in parentheses are the total numbers of genes in each ancestral mitochondrial genome. The white dots in the boxes of some *Mantamonas* proteins represent discrepancies in gene annotation between Moreira et al., (2024) and our analyses. Abbreviations of phyla: An, Ancyromonadida; Ma, Malawimonadida; Cm, CRuMs; Op, Opisthokonta; Am, Amoebozoa; Di, Discoba; Pr, Provora; Sa, SAR; Pa, Pancryptista; Ar, Archaeplastida.

More broadly, phylogenetic relationships within Opimoda+ including the phylogenetic position of Ancyromonadida are not conclusive (Brown et al., 2018; Eglit et al., 2024; Harada et al., 2024; Torruella et al., 2025; Williamson et al., 2025; Yazaki et al., 2025). Therefore, it is difficult to trace the reductive evolution of mitochondrial genome repertoires lineage by lineage within Opimoda+. Nevertheless, it is likely that the last common ancestor of Opimoda+ retained at least all of the 69 protein-coding genes detected in the mitochondrial genomes of Amorphea, CRuMs, Malawimonadida, and Ancyromonadida. If so, then six ribosomal protein-coding genes for *rps1*, *rps11*, *rpl18-20*, *rpl31*, and three cytochrome c maturase system I protein-coding genes, *ccmB*, *C*, and *F*, have very likely been lost within the Opimoda+ along the lineage leading to the Ancyromonadida; these genes appear to have been lost multiple times in eukaryotic evolution. Based on the same reasoning, the mitochondrial genome in LECA likely encoded at least 75 proteins, which are all of the protein-coding genes found in mitochondrial genomes. According to the recent phylogenomic analyses that estimated the root of the eukaryotes on the branch splitting Opimoda+ and Diphoda+ (Williamson et al., 2025), an ancestor of Opimoda+ had lost just four RNA polymerase subunit genes (*rpoA-D*), *rpl27* and *cox15*, on the descendant branch from LECA prior to diversification of Opimoda+. On the other side of the root, five ribosomal protein-coding genes for *rps5*, *rps6*, *rps16*, *rpl23*, and *rpl36*, and genes for sec translocator system except for *secY* were lost along the lineage from LECA to the common ancestor of extant Diphoda+. It appears that the vast majority of gene losses from the original alphaproteobacterial-related endosymbionts that gave rise to mitochondria were likely completed prior to the LECA (Richards et al., 2024; Roger et al., 2017) that might have possessed hundreds or thousands of protein-coding genes (Ettema and Andersson, 2009).

## CONCLUSIONS

In this study, we sequenced and analyzed the complete mitochondrial genomes of three Ancyromonadida species, revealing structural and genetic features that expand our understanding of mitochondrial genome evolution. We found that these genomes are circularly mapping molecules and contain PCIRs, which are patchily distributed across eukaryotes. Additionally, we identified *rps5*, a gene never detected in any mitochondrial genome. These findings provide new insights into the ancestral mitochondrial genome structure and gene repertoire. A more detailed picture of mitochondrial PCIR evolution and gene loss events along eukaryote phylogeny will be afforded by an improved understanding of the deep relationships within Opimoda+ as well as within Diphoda+ and on the availability of mitochondrial genomes from more diverse, deeply-diverging groups. Thus, we suggest that these inferences are revisited again whenever new mitochondrial genomes from diverse under-studied protists are determined.

## ACKNOWLEDGMENTS

We would like to thank Dr. Aaron A. Heiss at Kyungpook National University for providing the cell cultures. This work was in part supported by the Japan Society for Promotion of Sciences (JSPS) projects, 18J02091 (awarded to T. S.), 22KJ0401 (awarded to R. H.), 24K21929 (awarded to R. K.), 23K27226 and BPI06050 (awarded to Y. I.), a Discovery grant RGPIN-2022-05430 awarded to A. J. R. by the Natural Sciences and Engineering Research Council of Canada. R. H. was also supported by JSPS Overseas Research Fellowships. R. K. was also supported by a grant from Institute for Fermentation, Osaka (G-2024-1-011). A. Y. was also supported by the World Premier International Research Center Initiative (WPI Initiative), MEXT, Japan.

## Data availability Statement

The five mitochondrial genomes sequenced in this study were deposited in DDBJ database under accession nos. LC835033-5, LC842150 and LC860085. The protein tertiary structures predicted by AlphaFold3 and output files from InterProScan and Foldseek are available on Zenodo (https://doi.org/10.5281/zenodo.15079085).

## SUPPORTING INFORMATION

**Table S1.** Annotation results of uncharacterized proteins. All additional annotation results for ORFs which were not confidently annotated by Mfannot and PSI-BLAST were shown. Hits with significant scores were highlighted in yellow.

**FIGURE S1.** IGV screenshots showing DNA read mapping results. The lower left circle shows the four boundaries between inverted repeats (arrows) and single-copy regions. Each IGV screenshot shows 100 bp upstream and 100 bp downstream of the boundary. For each of the four Ancyromonadida genomes, the boundaries were confirmed to be continuously covered by DNA reads.

## REFERENCES

Abrahamsen M. S., Templeton T. J., Enomoto S., Abrahante J. E., Zhu G., Lancto C. A., Deng M., Liu C., Widmer G., Tzipori S., Buck G. A., Xu P., Bankier A. T., Dear P. H., Konfortov B. A., Spriggs H. F., Iyer L., Anantharaman V., Aravind L. & Kapur V. (2004). Complete Genome Sequence of the Apicomplexan, *Cryptosporidium parvum*. Science, 304(5669), 441–445. Available from: 10.1126/science.1094786

Abramson J., Adler J., Dunger J., Evans R., Green T., Pritzel A., Ronneberger O., Willmore L., Ballard A. J., Bambrick J., Bodenstein S. W., Evans D. A., Hung C.-C., O’Neill M., Reiman D., Tunyasuvunakool K., Wu Z., Žemgulytė A., Arvaniti E., Beattie C., Bertolli O., Bridgland A., Cherepanov A., Congreve M., Cowen-Rivers A. I., Cowie A., Figurnov M., Fuchs F. B., Gladman H., Jain R., Khan Y. A., Low C. M. R., Perlin K., Potapenko A., Savy P., Singh S., Stecula A., Thillaisundaram A., Tong C., Yakneen S., Zhong E. D., Zielinski M., Žídek A., Bapst V., Kohli P., Jaderberg M., Hassabis D. & Jumper J. M. (2024). Accurate structure prediction of biomolecular interactions with AlphaFold 3. Nature, 630(8016), 493–500. Available from: 10.1038/s41586-024-07487-w

Altschul S. F., Madden T. L., Schäffer A. A., Zhang J., Zhang Z., Miller W. & Lipman D. J. (1997). Gapped BLAST and PSI-BLAST: a new generation of protein database search programs. Nucleic Acids Research, 25(17), 3389–3402. Available from: 10.1093/nar/25.17.3389

Bolger A. M., Lohse M. & Usadel B. (2014). Trimmomatic: a flexible trimmer for Illumina sequence data. Bioinformatics, 30(15), 2114–2120. Available from: 10.1093/bioinformatics/btu170

Brown M. W., Heiss A. A., Kamikawa R., Inagaki Y., Yabuki A., Tice A. K., Shiratori T., Ishida K., Hashimoto T., Simpson A. G. B. & Roger A. J. (2018). Phylogenomics places orphan protistan lineages in a novel eukaryotic super-group. Genome Biology and Evolution, 10(2), 427–433. Available from: 10.1093/gbe/evy014

Burger G., Gray M. W., Forget L. & Lang B. F. (2013). Strikingly Bacteria-Like and Gene-Rich Mitochondrial Genomes throughout Jakobid Protists. Genome Biology and Evolution, 5(2), 418–438. Available from: 10.1093/gbe/evt008

Burger G., Gray M. W. & Franz Lang B. (2003). Mitochondrial genomes: anything goes. Trends in Genetics, 19(12), 709–716. Available from: 10.1016/j.tig.2003.10.012

Camacho C., Coulouris G., Avagyan V., Ma N., Papadopoulos J., Bealer K. & Madden T. L. (2009). BLAST+: architecture and applications. BMC Bioinformatics, 10, 421. Available from: 10.1186/1471-2105-10-421

Chen S., Zhou Y., Chen Y. & Gu J. (2018). fastp: an ultra-fast all-in-one FASTQ preprocessor. Bioinformatics, 34(17), i884–i890. Available from: 10.1093/bioinformatics/bty560

Choi I.-S., Jansen R. & Ruhlman T. (2019). Lost and Found: Return of the Inverted Repeat in the Legume Clade Defined by Its Absence. Genome Biology and Evolution, 11(4), 1321–1333. Available from: 10.1093/gbe/evz076

Derelle R., Torruella G., Klimeš V., Brinkmann H., Kim E., Vlček Č., Lang B. F. & Eliáš M. (2015). Bacterial proteins pinpoint a single eukaryotic root. Proceedings of the National Academy of Sciences, 112(7), E693–E699. Available from: 10.1073/pnas.1420657112

Dobáková E., Flegontov P., Skalický T. & Lukeš J. (2015). Unexpectedly Streamlined Mitochondrial Genome of the Euglenozoan *Euglena gracilis*. Genome Biology and Evolution, 7(12), 3358–3367. Available from: 10.1093/gbe/evv229

Eglit Y., Shiratori T., Jerlström-Hultqvist J., Williamson K., Roger A. J., Ishida K. & Simpson A. G. B. (2024). *Meteora sporadica*, a protist with incredible cell architecture, is related to Hemimastigophora. Current Biology, 34(2), 451–459.e6. Available from: 10.1016/j.cub.2023.12.032

Ettema T. J. G. & Andersson S. G. E. (2009). The α-proteobacteria: the Darwin finches of the bacterial world. Biology Letters, 5(3), 429–432. Available from: 10.1098/rsbl.2008.0793

Feagin J. E. (2000). Mitochondrial genome diversity in parasites. International Journal for Parasitology, 30(4), 371–390. Available from: 10.1016/S0020-7519(99)00190-3

Gastineau R., Harðardóttir S., Guilmette C., Lemieux C., Turmel M., Otis C., Boyle B., Levesque R. C., Gauthier J., Potvin M. & Lovejoy C. (2023). Mitochondrial genome sequence of the protist Ancyromonas sigmoides Kent, 1881 (Ancyromonadida) from the Sugluk Inlet, Hudson Strait, Nunavik, Québec. Front. Microbiol., 14, 1275665. Available from: 10.3389/fmicb.2023.1275665

Glücksman E., Snell E. A. & Cavalier-Smith T. (2013). Phylogeny and evolution of Planomonadida (Sulcozoa): Eight new species and new genera *Fabomonas* and *Nutomonas*. European Journal of Protistology, 49(2), 179–200. Available from: 10.1016/j.ejop.2012.08.007

Gray M. W., Lang B. F., Cedergren R., Golding G. B., Lemieux C., Sankoff D., Turmel M., Brossard N., Delage E., Littlejohn T. G., Plante I., Rioux P., Saint-Louis D., Zhu Y. & Burger G. (1998). Genome structure and gene content in protist mitochondrial DNAs. Nucleic Acids Research, 26(4), 865–878. Available from: 10.1093/nar/26.4.865

Greiner S., Lehwark P. & Bock R. (2019). OrganellarGenomeDRAW (OGDRAW) version 1.3.1: expanded toolkit for the graphical visualization of organellar genomes. Nucleic Acids Research, 47(W1), W59–W64. Available from: 10.1093/nar/gkz238

Hallgren J., Tsirigos K. D., Pedersen M. D., Armenteros J. J. A., Marcatili P., Nielsen H., Krogh A. & Winther O. (2022). DeepTMHMM predicts alpha and beta transmembrane proteins using deep neural networks. bioRxiv, 2022.04.08.487609. Available from: 10.1101/2022.04.08.487609

Harada R., Hirakawa Y., Yabuki A., Kim E., Yazaki E., Kamikawa R., Nakano K., Eliáš M. & Inagaki Y. (2024). Encyclopedia of family A DNA polymerases localized in organelles: evolutionary contribution of bacteria including the proto-mitochondrion. Molecular Biology and Evolution, 41(2), msae014. Available from: 10.1093/molbev/msae014

He D., Fu C.-J. & Baldauf S. L. (2016). Multiple Origins of Eukaryotic *cox15* Suggest Horizontal Gene Transfer from Bacteria to Jakobid Mitochondrial DNA. Molecular Biology and Evolution, 33(1), 122–133. Available from: 10.1093/molbev/msv201

Heiss A. A., Walker G. & Simpson A. G. B. (2010). Clarifying the Taxonomic Identity of a Phylogenetically Important Group of Eukaryotes: *Planomonas* is a Junior Synonym of *Ancyromonas*. Journal of Eukaryotic Microbiology, 57(3), 285–293. Available from: 10.1111/j.1550-7408.2010.00477.x

Janouškovec J., Tikhonenkov D. V., Burki F., Howe A. T., Rohwer F. L., Mylnikov A. P. & Keeling P. J. (2017). A New Lineage of Eukaryotes Illuminates Early Mitochondrial Genome Reduction. Current Biology, 27(23), 3717–3724.e5. Available from: 10.1016/j.cub.2017.10.051

Janouškovec J., Tikhonenkov D. V., Mikhailov K. V., Simdyanov T. G., Aleoshin V. V., Mylnikov A. P. & Keeling P. J. (2013). Colponemids Represent Multiple Ancient Alveolate Lineages. Current Biology, 23(24), 2546–2552. Available from: 10.1016/j.cub.2013.10.062

Jones P., Binns D., Chang H.-Y., Fraser M., Li W., McAnulla C., McWilliam H., Maslen J., Mitchell A., Nuka G., Pesseat S., Quinn A. F., Sangrador-Vegas A., Scheremetjew M., Yong S.-Y., Lopez R. & Hunter S. (2014). InterProScan 5: genome-scale protein function classification. Bioinformatics, 30(9), 1236–1240. Available from: 10.1093/bioinformatics/btu031

Käll L., Krogh A. & Sonnhammer E. L. L. (2004). A Combined Transmembrane Topology and Signal Peptide Prediction Method. Journal of Molecular Biology, 338(5), 1027–1036. Available from: 10.1016/j.jmb.2004.03.016

Kamikawa R., Inagaki Y. & Sako Y. (2007). Fragmentation of Mitochondrial Large Subunit rRNA in the Dinoflagellate *Alexandrium catenella* and the Evolution of rRNA structure in Alveolate Mitochondria. Protist, 158(2), 239–245. Available from: 10.1016/j.protis.2006.12.002

Kamikawa R., Shiratori T., Ishida K., Miyashita H. & Roger A. J. (2016). Group II Intron-Mediated *Trans*-Splicing in the Gene-Rich Mitochondrial Genome of an Enigmatic Eukaryote, *Diphylleia rotans*. Genome Biology and Evolution, 8(2), 458–466. Available from: 10.1093/gbe/evw011

Kamikawa R., Tanifuji G., Kawachi M., Miyashita H., Hashimoto T. & Inagaki Y. (2015). Plastid Genome-Based Phylogeny Pinpointed the Origin of the Green-Colored Plastid in the Dinoflagellate *Lepidodinium chlorophorum*. Genome Biology and Evolution, 7(4), 1133–1140. Available from: 10.1093/gbe/evv060

Kaur B., Záhonová K., Valach M., Faktorová D., Prokopchuk G., Burger G. & Lukeš J. (2020). Gene fragmentation and RNA editing without borders: eccentric mitochondrial genomes of diplonemids. Nucleic Acids Research, 48(5), 2694–2708. Available from: 10.1093/nar/gkz1215

Kayama M., Maciszewski K., Yabuki A., Miyashita H., Karnkowska A. & Kamikawa R. (2020). Highly Reduced Plastid Genomes of the Non-photosynthetic Dictyochophyceans *Pteridomonas* spp. (Ochrophyta, SAR) Are Retained for tRNA-Glu-Based Organellar Heme Biosynthesis. Front. Plant Sci., 11, 602455. Available from: 10.3389/fpls.2020.602455

van Kempen M., Kim S. S., Tumescheit C., Mirdita M., Lee J., Gilchrist C. L. M., Söding J. & Steinegger M. (2024). Fast and accurate protein structure search with Foldseek. Nat Biotechnol, 42(2), 243–246. Available from: 10.1038/s41587-023-01773-0

Kim D., Paggi J. M., Park C., Bennett C. & Salzberg S. L. (2019). Graph-based genome alignment and genotyping with HISAT2 and HISAT-genotype. Nat Biotechnol, 37(8), 907–915. Available from: 10.1038/s41587-019-0201-4

Krämer C., Boehm C. R., Liu J., Ting M. K. Y., Hertle A. P., Forner J., Ruf S., Schöttler M. A., Zoschke R. & Bock R. (2024). Removal of the large inverted repeat from the plastid genome reveals gene dosage effects and leads to increased genome copy number. Nat. Plants, 10(6), 923–935. Available from: 10.1038/s41477-024-01709-9

Krogh A., Larsson B., von Heijne G. & Sonnhammer E. L. L. (2001). Predicting transmembrane protein topology with a hidden Markov model: application to complete genomes. Journal of Molecular Biology, 305(3), 567–580. Available from: 10.1006/jmbi.2000.4315

Lang B. F., Beck N., Prince S., Sarrasin M., Rioux P. & Burger G. (2023). Mitochondrial genome annotation with MFannot: a critical analysis of gene identification and gene model prediction. Front. Plant Sci., 14, 1222186. Available from: 10.3389/fpls.2023.1222186

Leger M. M., Kolisko M., Kamikawa R., Stairs C. W., Kume K., Čepička I., Silberman J. D., Andersson J. O., Xu F., Yabuki A., Eme L., Zhang Q., Takishita K., Inagaki Y., Simpson A. G. B., Hashimoto T. & Roger A. J. (2017). Organelles that illuminate the origins of *Trichomonas* hydrogenosomes and *Giardia* mitosomes. Nat Ecol Evol, 1(4), 1–7. Available from: 10.1038/s41559-017-0092

Lonergan K. M. & Gray M. W. (1996). Expression of a Continuous Open Reading Frame Encoding Subunits 1 and 2 of Cytochrome *c* Oxidase in the Mitochondrial DNA of *Acanthamoeba castellanii*. Journal of Molecular Biology, 257(5), 1019–1030. Available from: 10.1006/jmbi.1996.0220

Martin M. (2011). Cutadapt removes adapter sequences from high-throughput sequencing reads. EMBnet.journal, 17(1), 10–12. Available from: 10.14806/ej.17.1.200

Matsuo E., Morita K., Nakayama T., Yazaki E., Sarai C., Takahashi K., Iwataki M. & Inagaki Y. (2022). Comparative Plastid Genomics of Green-Colored Dinoflagellates Unveils Parallel Genome Compaction and RNA Editing. Front. Plant Sci., 13, 918543. Available from: 10.3389/fpls.2022.918543

Moreira D., Blaz J., Kim E. & Eme L. (2024). A gene-rich mitochondrion with a unique ancestral protein transport system. Current Biology, 34(16), 3812–3819.e3. Available from: 10.1016/j.cub.2024.07.017

Nawrocki E. P. & Eddy S. R. (2013). Infernal 1.1: 100-fold faster RNA homology searches. Bioinformatics, 29(22), 2933–2935. Available from: 10.1093/bioinformatics/btt509

Nishimura Y., Kume K., Sonehara K., Tanifuji G., Shiratori T., Ishida K., Hashimoto T., Inagaki Y. & Ohkuma M. (2020). Mitochondrial Genomes of *Hemiarma marina* and *Leucocryptos marina* Revised the Evolution of Cytochrome *c* Maturation in Cryptista. Front. Ecol. Evol., 8, 140. Available from: 10.3389/fevo.2020.00140

Nishimura Y., Tanifuji G., Kamikawa R., Yabuki A., Hashimoto T. & Inagaki Y. (2016). Mitochondrial Genome of *Palpitomonas bilix*: Derived Genome Structure and Ancestral System for Cytochrome *c* Maturation. Genome Biology and Evolution, 8(10), 3090–3098. Available from: 10.1093/gbe/evw217

Norman J. E. & Gray M. W. (2001). A Complex Organization of the Gene Encoding Cytochrome Oxidase Subunit 1 in the Mitochondrial Genome of the Dinoflagellate, Crypthecodinium cohnii: Homologous Recombination Generates Two Different cox1 Open Reading Frames. J Mol Evol, 53(4), 351–363. Available from: 10.1007/s002390010225

Nurk S., Meleshko D., Korobeynikov A. & Pevzner P. A. (2017). metaSPAdes: a new versatile metagenomic assembler. Genome Res., 27(5), 824–834. Available from: 10.1101/gr.213959.116

Oborník M. & Lukeš J. (2015). The Organellar Genomes of *Chromera* and *Vitrella*, the Phototrophic Relatives of Apicomplexan Parasites. Annual Review of Microbiology, 69, 129–144. Available from: 10.1146/annurev-micro-091014-104449

Palmer J. D. & Thompson W. F. (1982). Chloroplast DNA rearrangements are more frequent when a large inverted repeat sequence is lost. Cell, 29(2), 537–550. Available from: 10.1016/0092-8674(82)90170-2

Pérez-Brocal V., Shahar-Golan R. & Clark C. G. (2010). A Linear Molecule with Two Large Inverted Repeats: The Mitochondrial Genome of the Stramenopile *Proteromonas lacertae*. Genome Biology and Evolution, 2, 257–266. Available from: 10.1093/gbe/evq015

Richards T. A., Eme L., Archibald J. M., Leonard G., Coelho S. M., Mendoza A. de, Dessimoz C., Dolezal P., Fritz-Laylin L. K., Gabaldón T., Hampl V., Kops G. J. P. L., Leger M. M., Lopez-Garcia P., McInerney J. O., Moreira D., Muñoz-Gómez S. A., Richter D. J., Ruiz-Trillo I., Santoro A. E., Sebé-Pedrós A., Snel B., Stairs C. W., Tromer E. C., Hooff J. J. E. van, Wickstead B., Williams T. A., Roger A. J., Dacks J. B. & Wideman J. G. (2024). Reconstructing the last common ancestor of all eukaryotes. PLOS Biology, 22(11), e3002917. Available from: 10.1371/journal.pbio.3002917

Roger A. J., Muñoz-Gómez S. A. & Kamikawa R. (2017). The origin and diversification of mitochondria. Current Biology, 27(21), R1177–R1192. Available from: 10.1016/j.cub.2017.09.015

Slamovits C. H., Saldarriaga J. F., Larocque A. & Keeling P. J. (2007). The Highly Reduced and Fragmented Mitochondrial Genome of the Early-branching Dinoflagellate *Oxyrrhis marina* Shares Characteristics with both Apicomplexan and Dinoflagellate Mitochondrial Genomes. Journal of Molecular Biology, 372(2), 356–368. Available from: 10.1016/j.jmb.2007.06.085

Spencer D. F. & Gray M. W. (2011). Ribosomal RNA genes in *Euglena gracilis* mitochondrial DNA: fragmented genes in a seemingly fragmented genome. Mol Genet Genomics, 285(1), 19–31. Available from: 10.1007/s00438-010-0585-9

Tikhonenkov D. V., Janouškovec J., Mylnikov A. P., Mikhailov K. V., Simdyanov T. G., Aleoshin V. V. & Keeling P. J. (2014). Description of *Colponema vietnamica* sp.n. and *Acavomonas peruviana* n. gen. n. sp., Two New Alveolate Phyla (Colponemidia nom. nov. and Acavomonidia nom. nov.) and Their Contributions to Reconstructing the Ancestral State of Alveolates and Eukaryotes. PLOS ONE, 9(4), e95467. Available from: 10.1371/journal.pone.0095467

Tikhonenkov D. V., Mikhailov K. V., Gawryluk R. M. R., Belyaev A. O., Mathur V., Karpov S. A., Zagumyonnyi D. G., Borodina A. S., Prokina K. I., Mylnikov A. P., Aleoshin V. V. & Keeling P. J. (2022). Microbial predators form a new supergroup of eukaryotes. Nature, 612(7941), 714–719. Available from: 10.1038/s41586-022-05511-5

Torruella G., Galindo L. J., Moreira D. & López-García P. (2025). Phylogenomics of neglected flagellated protists supports a revised eukaryotic tree of life. Current Biology, 35(1), 198–207.e4. Available from: 10.1016/j.cub.2024.10.075

Valach M., Burger G., Gray M. W. & Lang B. F. (2014). Widespread occurrence of organelle genome-encoded 5S rRNAs including permuted molecules. Nucleic Acids Research, 42(22), 13764–13777. Available from: 10.1093/nar/gku1266

Verner Z., Basu S., Benz C., Dixit S., Dobáková E., Faktorová D., Hashimi H., Horáková E., Huang Z., Paris Z., Peña-Diaz P., Ridlon L., Týč J., Wildridge D., Zíková A. & Lukeš J. (2015). Malleable Mitochondrion of *Trypanosoma brucei*. *In*: Jeon K. W. (ed.), International Review of Cell and Molecular Biology. Vol. 315. Academic Press. p. 73–151. Available from: 10.1016/bs.ircmb.2014.11.001

Vosseberg J., van Hooff J. J. E., Köstlbacher S., Panagiotou K., Tamarit D. & Ettema T. J. G. (2024). The emerging view on the origin and early evolution of eukaryotic cells. Nature, 633(8029), 295–305. Available from: 10.1038/s41586-024-07677-6

Williams B. A. P., Hirt R. P., Lucocq J. M. & Embley T. M. (2002). A mitochondrial remnant in the microsporidian *Trachipleistophora hominis*. Nature, 418(6900), 865–869. Available from: 10.1038/nature00949

Williamson K., Eme L., Baños H., McCarthy C. G. P., Susko E., Kamikawa R., Orr R. J. S., Muñoz-Gómez S. A., Minh B. Q., Simpson A. G. B. & Roger A. J. (2025). A robustly rooted tree of eukaryotes reveals their excavate ancestry. Nature, 1–8. Available from: 10.1038/s41586-025-08709-5

Yang J., Harding T., Kamikawa R., Simpson A. G. B. & Roger A. J. (2017). Mitochondrial Genome Evolution and a Novel RNA Editing System in Deep-Branching Heteroloboseids. Genome Biology and Evolution, 9(5), 1161–1174. Available from: 10.1093/gbe/evx086

Yazaki E., Harada R., Isogai R., Bamba K., Ishida K., Inagaki Y. & Shiratori T. (2025). *Glissandra oviformis* n. sp.: a novel predatory flagellate illuminates the character evolution within the eukaryotic clade CRuMs. bioRxiv, 2025.02.18.638917. Available from: 10.1101/2025.02.18.638917

Yazaki E., Yabuki A., Nishimura Y., Shiratori T., Hashimoto T. & Inagaki Y. (2022). *Microheliella maris* possesses the most gene-rich mitochondrial genome in Diaphoretickes. Front. Ecol. Evol., 10. Available from: 10.3389/fevo.2022.1030570

Yubuki N., Torruella G., Galindo L. J., Heiss A. A., Ciobanu M. C., Shiratori T., Ishida K., Blaz J., Kim E., Moreira D., López-García P. & Eme L. (2023). Molecular and morphological characterization of four new ancyromonad genera and proposal for an updated taxonomy of the Ancyromonadida. Journal of Eukaryotic Microbiology, 70(6), e12997. Available from: 10.1111/jeu.12997

Zerbino D. R. & Birney E. (2008). Velvet: Algorithms for de novo short read assembly using de Bruijn graphs. Genome Res., 18(5), 821–829. Available from: 10.1101/gr.074492.107

Zhang Y. & Skolnick J. (2005). TM-align: a protein structure alignment algorithm based on the TM-score. Nucleic Acids Research, 33(7), 2302–2309. Available from: 10.1093/nar/gki524

