## Supplemental Figure 1 for "Complete mitochondrial genomes of ancyromonads provide clues for the gene content and genome structures of ancestral mitochondria"

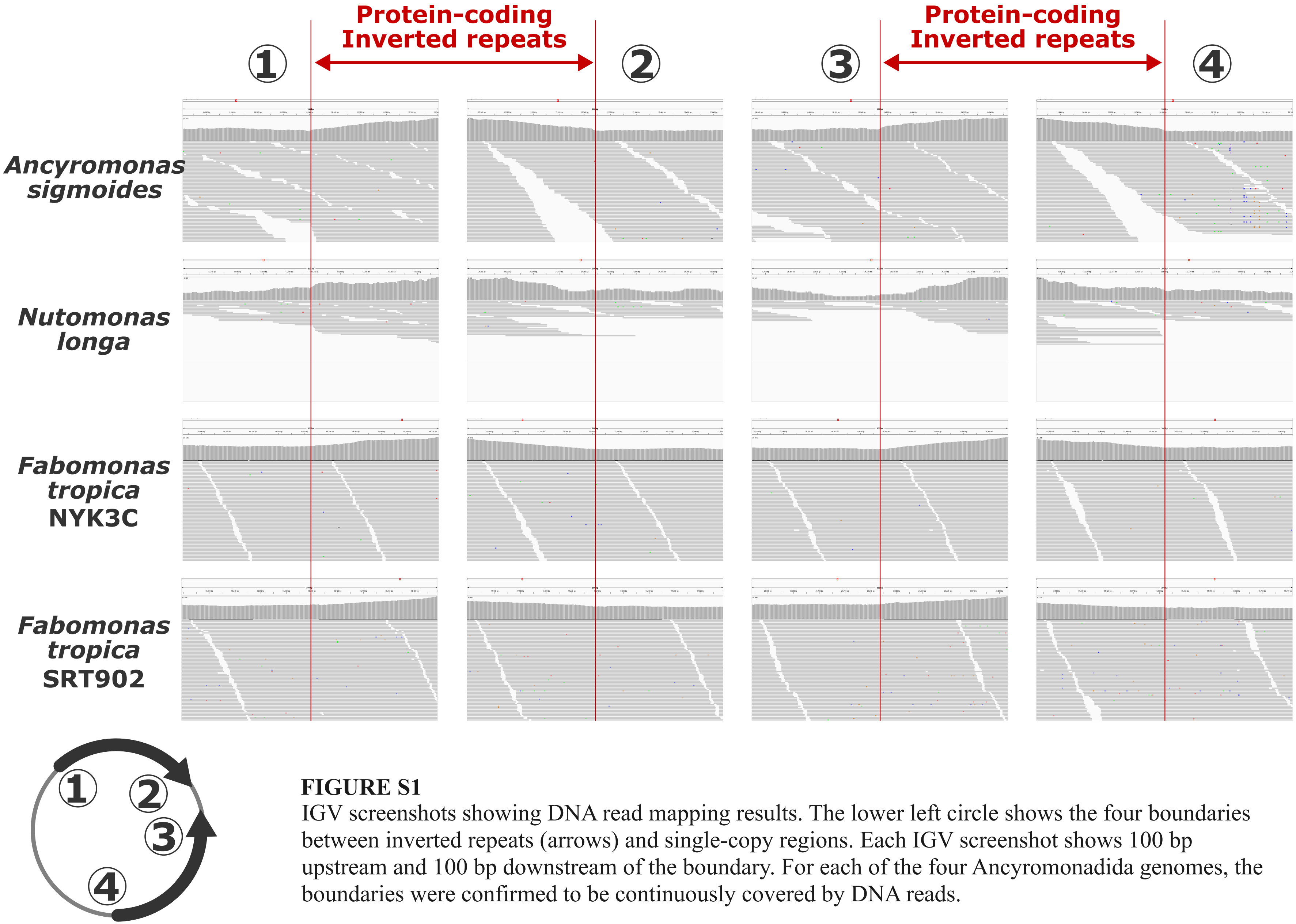


**FIGURE S1**
IGV screenshots showing DNA read mapping results. The lower left circle shows the four boundaries between inverted repeats (arrows) and single-copy regions. Each IGV screenshot shows 100 bp upstream and 100 bp downstream of the boundary. For each of the four Ancyromonadida genomes, the boundaries were confirmed to be continuously covered by DNA reads.
